# Endoglycosidase S enables a highly simplified clinical chemistry assay procedure for direct assessment of serum IgG undergalactosylation in chronic inflammatory disease

**DOI:** 10.1101/269795

**Authors:** Dieter Vanderschaeghe, Leander Meuris, Tom Raes, Hendrik Grootaert, Annelies Van Hecke, Xavier Verhelst, Frederique Van de Velde, Bruno Lapauw, Hans Van Vlierberghe, Nico Callewaert

## Abstract

Over the past 30 years, it has been firmly established that a wide spectrum of (autoimmune) diseases such as rheumatoid arthritis, Crohn’s and lupus, but also other pathologies like alcoholic and non-alcoholic steatohepatitis (ASH and NASH) are driven by chronic inflammation and are hallmarked by a reduced level of serum IgG galactosylation. IgG (under)galactosylation is a promising biomarker to assess disease severity, and monitor and adjust therapy. However, this biomarker has not been implemented in routine clinical chemistry due to a complex analytical procedure that necessitates IgG purification, which is difficult to perform and validate at high throughput.

We addressed this issue by using endo-β-N-acetylglucosaminidase from *Streptococcus pyogenes* (endoS) to specifically release Fc N-glycans in whole serum. The entire assay can be completed in a few hours and only entails adding endoS and labeling the glycans with APTS. Glycans are then readily analyzed through capillary electrophoresis. We demonstrate in two independent patient cohorts that IgG undergalactosylation levels obtained with this assay correlate very well with scores calculated from PNGaseF-released glycans of purified antibodies. Our new assay allows to directly and specifically measure the degree of IgG galactosylation in serum through a fast and completely liquid phase protocol, without the requirement for antibody purification. This should help advancing this biomarker towards clinical implementation.

## Introduction

IgG glycosylation has been studied for more than 30 years, revealing how glycan composition is altered in patients suffering from chronic inflammatory diseases. Early studies showed that β-1,4-galactosylation on IgG glycans is lowered in individuals with rheumatoid arthritis (RA) or primary osteoarthritis (1,2). Since then, other (chronic) inflammatory diseases have been shown to be associated with IgG undergalactosylation, including several cancers (3,4), inflammatory bowel disease (5,6), systemic lupus erythematosus (7) and liver disease (8–10). Remarkably, in RA and hepatitis B virus (HBV) patients, the glycosylation patterns normalize after treatment (11–13). IgG galactosylation can also differentiate between benign non-alcoholic fatty liver disease (NAFLD) and non-alcoholic steatohepatitis (NASH), a rapidly emerging disease (10). Many drugs for treatment of chronic inflammatory diseases are very expensive (e.g. anti-TNF biologicals) and come with significant side effects. A biomarker for low grade chronic systemic inflammation and objective treatment monitoring would therefore contribute to improved clinical practice.

Despite numerous publications presenting IgG undergalactosylation as a promising biomarker, current clinical tests for inflammation are largely limited to measuring C-reactive protein (CRP) in serum or to historical tests such as the erythrocyte sedimentation rate (ESR). CRP levels fluctuate on a daily basis and it is therefore not a suitable marker for measuring chronic disease activity. ESR is a cheap test that is routinely conducted at high throughput, but requires fresh blood and is confounded by several interfering factors (14).

Analyzing IgG glycosylation has so far not been feasible in routine clinical chemistry due to the complex and time consuming procedure that is required. Purifying the antibodies from patients’ sera is a prerequisite and is then usually followed by a denaturation step and PNGaseF deglycosylation to assess the sample N-glycosylation profile by MS, ultra-performance liquid chromatography (UPLC) or capillary electrophoresis (CE) (15–17). The mandatory purification step is difficult to implement in routine testing and validation of antibody purity (which can affect the outcome of the test) at high throughput is cumbersome. Attempts to facilitate the purification process have recently resulted in an automated high-throughput IgG purification and N-glycan sample preparation platform (18). Although promising, sample turnaround for 96 samples takes at least 30 hours and requires an expensive robotic liquid handling and UPLC system, limiting its potential for clinical implementation.

In 2001, Collin and Olsén (19) described an endo-β-N-acetylglucosaminidase secreted by the human pathogen *Streptococcus pyogenes*, designated endoS. EndoS likely suppresses host IgG effector functions, as it can deglycosylate native IgG. It hydrolyzes the N-glycan chitobiose core, leaving a single GlcNAc residue on the IgG heavy chain (20). The enzyme reportedly requires a natively folded IgG for its activity (21) and only cleaves Fc-linked glycans while leaving N-glycans on Fab and other proteins intact (22). Although it is generally accepted that endoS is specific for IgG Fc, activity towards HIV gp120 has been reported (23). Moreover, endoS specificity has never been tested in a complex mixture such as serum. Most Fc N-glycans are cleaved by endoS, although glycans with a bisecting GlcNAc have been shown to be resistant (24). The extent to which endoS-released glycans of patients’ sera resemble the total N-glycan profile of the corresponding purified antibodies remains unexplored.

Here we describe a simple and fast, liquid phase, endoS-based assay to specifically measure IgG galactosylation without the need for antibody purification, which overcomes the main bottleneck for widespread clinical use of this biomarker.

## Materials and Methods

### Materials

EndoS (IgGZERO™, Genovis) was reconstituted in ultrapure water at 20 U/µl. PNGaseF from *Flavobacterium meningosepticum* and sialidase from *Arthrobacter ureafaciens* were recombinantly produced as described previously (25). IgG, IgA and IgM from human serum (purity > 95%, Sigma-Aldrich) were reconstituted in PBS at 10 mg/ml (hereafter named ‘commercial IgG, IgA and IgM’).

### Patients

Serum samples from two independent cohorts (10,26) (*n*_*total*_ = 188) in which patients were enlisted for bariatric surgery, were obtained from an outpatient clinic (Ghent University Hospital) and stored at −80°C. Informed consent was given by all patients and the protocol was approved by the Hospital’s Ethics Committee.

### IgG purification and serum IgG depletion

IgG from 30 µl aliquots of sera was purified using a Protein G Spin Plate for IgG Screening (Thermo Scientific) according to the manufacturer’s instructions. Samples were eluted in 420 µl and concentrated/buffer exchanged to PBS with 100 kDa cut-off Amicon^®^ Ultra Centrifugal Filter Devices (Merck).

IgG-depleted sera were prepared with the ProteoPrep^®^ Immunoaffinity Albumin and IgG Depletion Kit (Sigma-Aldrich) according to the manufacturer’s instructions. Depleted sera were concentrated/buffer-exchanged to PBS with 10 kDa cut-off Amicon^®^ Centrifugal Filter devices to obtain original serum protein concentrations.

### Release of N-glycans

N-glycans were released with endoS or PNGaseF. For endoS catalyzed release, 2.5 µl of serum or purified IgG was incubated with 25 U of endoS for one hour at 37°C in a total volume of 10 µl in 150 mmol/L NaCl and 50 mmol/L Tris-HCl pH 8.5.

For PNGaseF catalyzed N-glycan release, 3 µl of serum or purified IgG was denatured by incubation at 95°C for 5 minutes after adding 2 µl 50 mmol/L Tris-HCl pH 8.5 containing 3.5% SDS. Samples were cooled and incubated for 1 hour at 37°C with 5 µl of deglycosylation mix, containing 11 mU PNGaseF in 50 mmol/L Tris-HCl pH 8.5 and 2% NP40.

### IgG specificity of endoS

To determine the protein specificity of endoS, serum and IgG-depleted serum were incubated with 25 U of endoS for 1 hour at 37°C in buffers with a pH ranging from 5.0 to 9.0: 50 mmol/L NH_4_Ac (pH 5.0), MES (pH 6.0), sodium phosphate (pH 7.4) or Tris-Cl (pH 8.0, 8.5 and 9.0), each containing 150 mmol/L NaCl. IgA from patient samples was depleted/purified by diluting 500 µl plasma into 2 ml of PBS, followed by centrifugation at 16,000 x g and supernatant filtering (0.22 µM). The sample was then applied three times over 0.5 ml of PBS-equilibrated peptide M agarose (Invivogen). The flowthrough after the third pass was collected as IgA-depleted plasma. The column was washed with 20 CV’s of PBS and IgA eluted with 2.5 ml of 200 mmol/L glycine pH 2.5 and neutralized with 2.5 ml of 1 M Tris-Cl pH 8.5. IgA and IgA-depleted plasma were buffer exchanged to 500 µl PBS with 3 and 30 kDa Amicon^®^ centrifugal filter devices respectively.

To determine whether endoS hydrolyses IgA or IgM glycans, 2.5 µl samples of commercial IgA, IgM, IgG, patient IgA and IgA-depleted plasma were digested with 25 U of endoS for 1 hour at 37°C in a total volume of 10 µl in 50 mmol/L NH_4_Ac pH 5.0 or 50 mmol/L Tris-HCl pH 8.5, each with 150 mmol/L NaCl.

### Labelling and CE analysis

Released glycans were fluorescently labelled with APTS and analyzed by CE. Sample preparation, exoglycosidase digests (optional) and electrophoresis were performed as previously described (27). Briefly, 5 µl of labelling solution was added to an equal volume of crude digest or dry samples, followed by overnight incubation at 37°C. Excess label was removed by size-exclusion chromatography. Labelled N-glycans were then optionally desialylated, followed by CE on an ABI3130 DNA sequencer.

Glycan symbolic representations are according to the Consortium for Functional Glycomics guidelines (28). Glycan structures, nomenclature and symbols are summarized in Supplemental Fig. 1.

**Figure 1:**
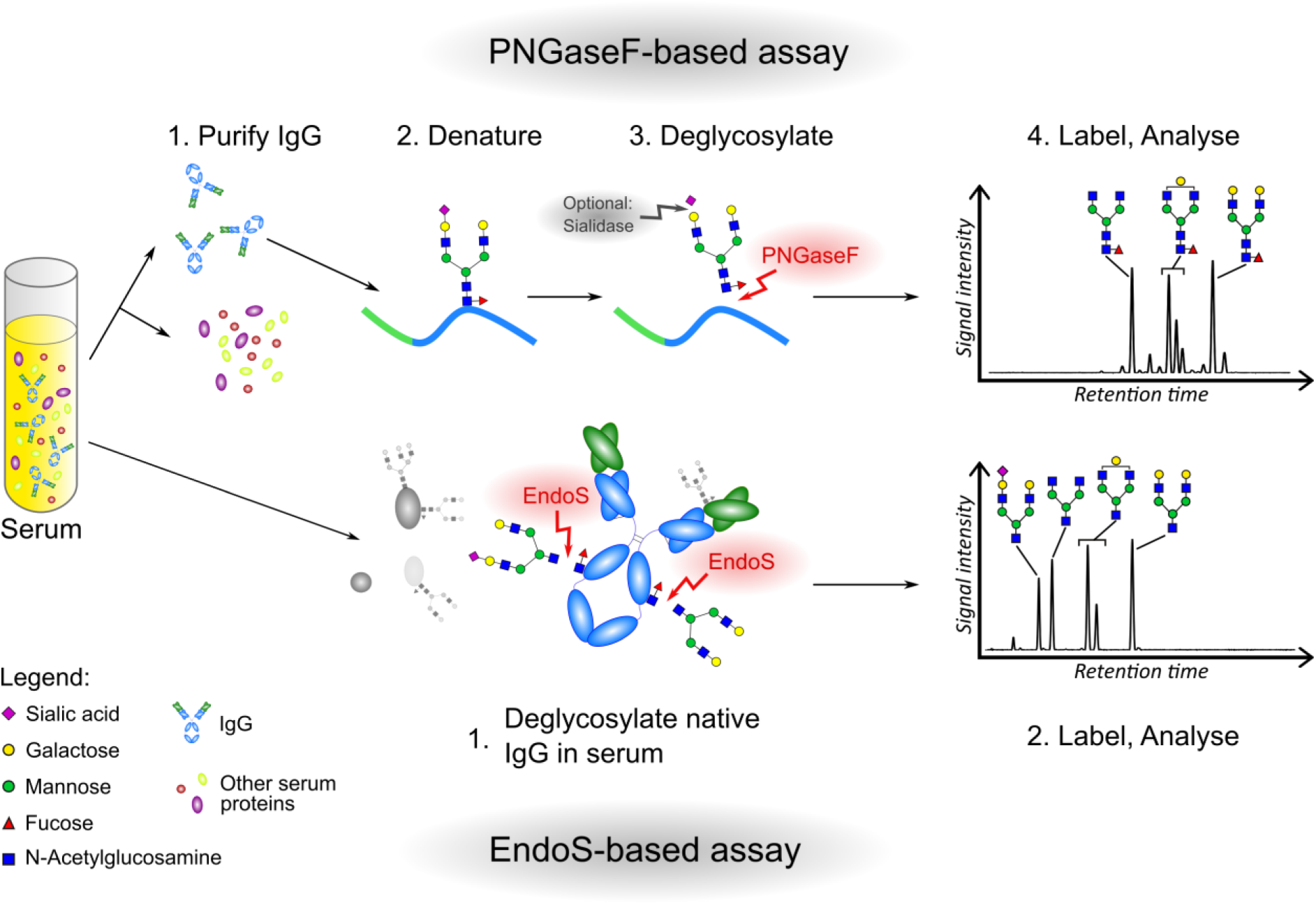
Overview of the assay. In traditional PNGaseF-based assays to assess IgG Fc glycan galactosylation levels, IgG has to be purified from serum, denatured and then deglycosylated (and optionally desialylated) before the glycans can be analysed. We here propose a new assay, based on the use of endoS, in which the purification and denaturation steps can be omitted, as well as the use of sialidase. Due to endoS’ specificity for IgG Fc glycans on folded IgG, the resulting profile can be used to determine IgG Fc galactosylation levels.

### Calculations and statistics

The relevant peaks in the glycan profiles were quantified with GeneMapper^®^ v3.7 (Applied Biosystems). The level of galactosylation (undergalactosylation score, UGS) from PNGaseF profiles was calculated as the ratio of NGA2F (G0F) glycan over the total peak height (Supplemental Fig. 2: PNGase UGS). The UGS from endoS profiles was calculated as the sum of peak heights of non-galactosylated glycans normalized to total peak height, taking into account the number of antennae for each glycan (Supplemental Fig. 2: endoS UGS).

**Figure 2:**
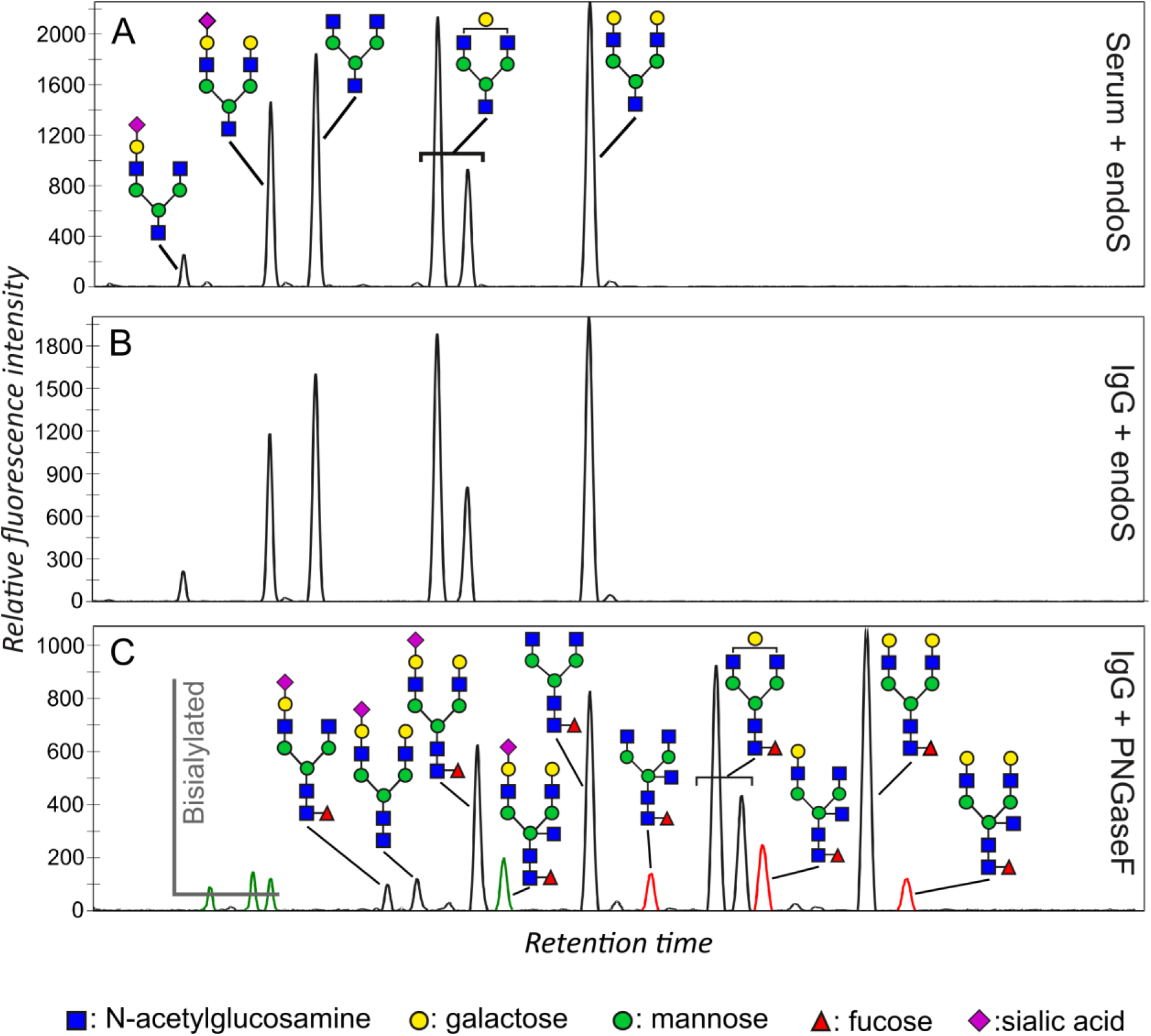
CE-LIF profiles of endoS and PNGaseF digested N-glycans. **A)** EndoS-released N-glycans from serum of a healthy individual. **B)** EndoS-released N-glycans from purified whole IgG of the same individual. **C)** PNGaseF-released glycans from purified whole IgG of the same individual. Peaks in red represent the bisecting GlcNAc-containing structures on the Fc fragment that are resistant to endoS hydrolysis. Bisialylated glycans and the bisected monosialylated N-glycan (indicated in green) are almost exclusively coming from Fab and were thus not hydrolysed by endoS (see Supplementary Figure 4 for additional data on the origin of the different glycan peaks).

Statistical analyses were performed with R v3.4. The compound patient cohort (n_total_ = 188) was used in all analyses. Correlations were addressed with Pearson’s r. Outliers from linear regression were defined as having studentized residuals higher than 3 or lower than −3. The glycan dataset can be found in the Supplemental spreadsheet.

## Results

### Assay and N-glycan profile characterization

To specifically analyze IgG N-glycosylation in serum using endoS, we developed a new method (Fig. 1), based on procedures for profiling serum N-glycosylation that we previously optimized (25). In contrast to PNGaseF, which is commonly used to remove N-glycans from denatured proteins (Fig. 1, top), endoS activity requires native (dimeric) IgG Fc fragments (Fig. 1, bottom). Only N-glycans from IgG Fc are expected to be released by endoS while leaving glycans on the Fab fragment and other serum glycoproteins unaffected, allowing the omission of IgG purification and denaturation steps.

To identify the glycan structures that are released by endoS, we analyzed endoS-released N-glycans from serum of a healthy individual and from IgG that was purified from the same serum sample (Fig. 2A and 2B). These CE profiles revealed six major glycan peaks. Purified IgG, again from the same serum sample, was also digested with PNGaseF for full N-glycan profiling (Fig. 2C). Comparing digests on serum and IgG showed that endoS not only cleaves the same N-glycans in both samples, but we also observed similar relative peak heights (Fig. 2A and 2B). These first results indicated the feasibility of using endoS in serum for specific analysis of IgG glycosylation.

Since endoS hydrolyses N-glycans in their chitobiose core, all the structures are truncated at the reducing end as compared to PNGaseF-released N-glycans, which hydrolyses the amide bond between the N-glycan and the asparagine side chain amine.

All endoS-released N-glycan structures were confirmed with exoglycosidase digests and subsequent CE analysis of the resulting fragments (Supplemental Fig. 3). The N-glycans were all biantennary complex type N-glycans that are typically present on IgG Fc. We found no structures with bisecting GlcNAc, although about 10% of IgG Fc glycans are modified with such a structure (29). This is consistent with earlier reports that endoS does not hydrolyse glycans with a bisecting GlcNAc (24).

**Figure 3:**
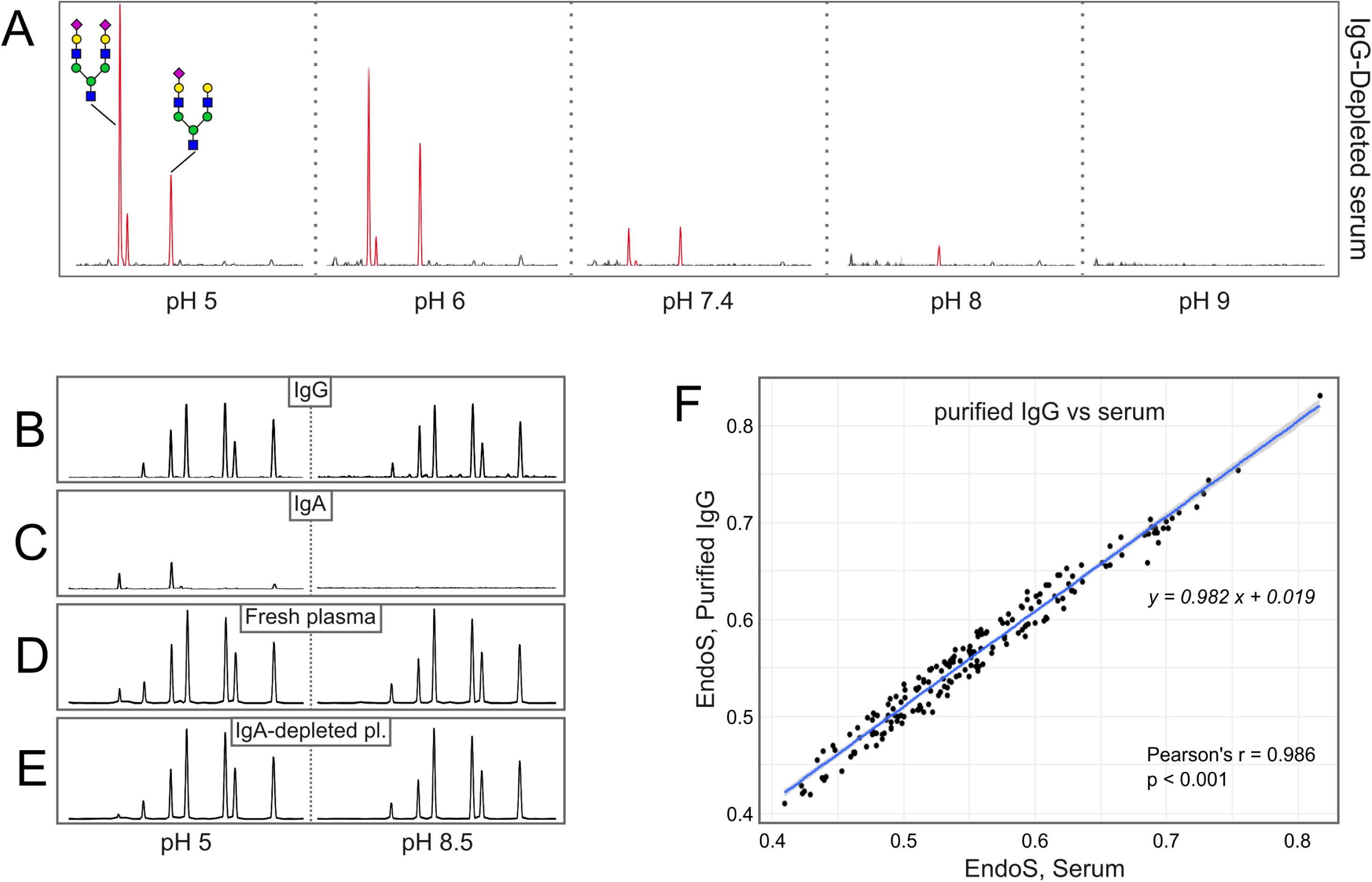
IgG specificity of EndoS in serum at different pH values. **A)** At pH values below 8.0, EndoS released bi- and monosialylated biantennary glycans from IgG-depleted serum samples. These glycans were not released from IgG-depleted serum samples at pH values above 8. **B)** The relative peak heights in the N-glycan profile of commercial IgG (purity > 95%) were independent of the assay buffer. **C)** Profiles of IgA, purified from patient samples. EndoS releases the same bi- and monosialylated biantennary N-glycans as found in IgG-depleted serum (Figure 3A) at pH 5.0. At pH 8.5, endoS does not hydrolyse any IgA glycans. **D)** Profiles of whole plasma from the same patient are pH dependent: at pH 8.5 a profile similar to the commercial IgG profile is observed. At pH 5.0, the profile contains IgG peaks as well as the same peaks as in the IgA profile and the depleted serum (Figure 3A and 3D). **E)** EndoS profiles of the same sample after IgA-depletion results similar profiles at either pH and also similar to the IgG profile. Note that 100 % IgA depletion is extremely difficult to achieve. Consequently, a small peak representing IgA-derived glycans can still be observed in the pH 5.0 profile of IgA-depleted samples. **F)** Serum and purified IgG from 188 patients and healthy controls were digested by endoS. The samples were labelled and analysed on an ABI3130 DNA sequencer. The UGS scores were calculated from the CE profiles. Scatterplots of undergalactosylation scores correlate very well for whole serum and IgG purified from the same serum samples, indicating that the presence of serum (glyco)proteins and other serum components do not change the assay outcome.

To identify endoS resistant IgG glycans, an endoS digest was performed on commercial whole IgG, followed by separation of the Fab and Fc fragments and analysis of the remaining glycans on both (Supplemental Fig. 4). This revealed that Fc glycans with a bisecting GlcNAc (peaks indicated in red in Fig. 2C) indeed remain intact after endoS treatment. Bisialylated N-glycans (peaks indicated in green in Fig. 2C) are not released by endoS, but are originating almost exclusively from the Fab fragments (Supplemental Fig. 4), which are no substrate for endoS. The lack of bisialylated N-glycans in the endoS profiles is consistent with reports that less than 0.5% of Fc glycans are bisialylated (29). Taken together, these observations support the view that, when using purified IgG as the substrate, endoS is specific for the Fc portion. Glycans with a bisecting GlcNAc, which constitute about 10% of Fc glycans, are resistant to endoS hydrolysis.

**Figure 4:**
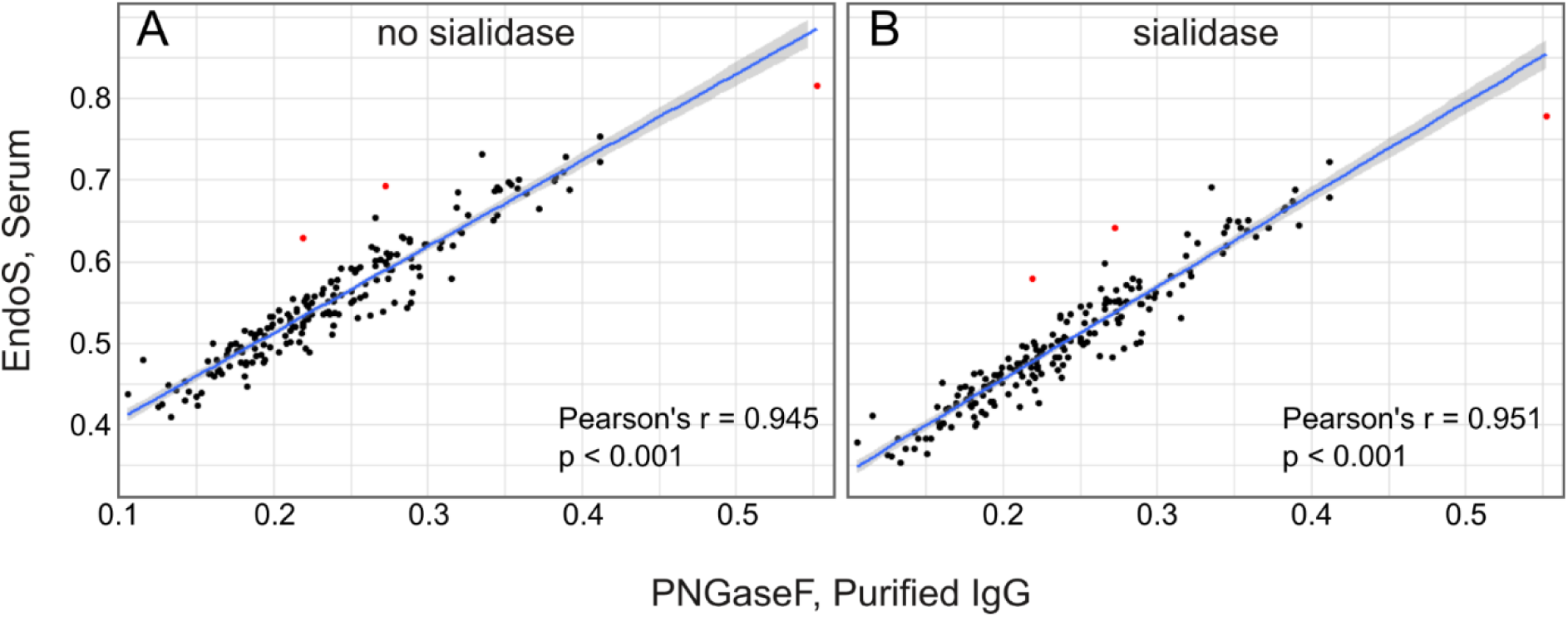
Correlation between galactosylation levels calculated from serum derived profiles versus purified IgG derived profiles. Serum and purified IgG from 188 patients and healthy controls were digested by endoS or PNGaseF. The samples were labelled and analysed on an ABI3130 DNA sequencer. The UGS scores were calculated from the CE profiles. Scatterplots of undergalactosylation scores are shown for endoS profiles of serum versus PNGaseF derived profiles for purified IgG from the same serum samples. **A)** undergalactosylation scores calculated from endoS-derived profiles versus PNGaseF-derived profiles. EndoS-derived glycans were not treated with sialidase. **B)** undergalactosylation scores calculated from endoS-derived profiles versus PNGaseF-derived profiles. EndoS-derived glycans were treated with sialidase prior to analysis. Pearson’s r correlation coefficients are shown in the right lower corner of all panels, along with p-values for each set. The linear regression lines (blue) with their 95% confidence intervals (grey) are also shown for each comparison. The three datapoints depicted in red were identified as outliers by studentized residuals > 3. Two out of these represent samples in which the IgG purification was suboptimal.

### IgG specificity of endoS in serum

To assess whether endoS cleaves only IgG N-glycans in serum, we performed endoS digests on IgG-depleted serum. We previously demonstrated that up to 99% of IgG is removed by this procedure, without affecting other serum proteins (30). Treating fresh serum (or plasma) with endoS in different pH conditions (pH 5.0 – 9.0), revealed the release of an increasing amount of sialylated N-glycans from proteins other than IgG with decreasing pH (Fig. 3A). Exoglycosidase digests showed these peaks to represent mono- and bisialylated, fully galactosylated biantennary complex type glycans. These glycans were not observed when using plasma or serum that had been freeze-thawed several times as the starting material (data not shown). Adding a physiological amount of commercial IgG (10 µg/µL) back to IgG-depleted serum shows that non-IgG N-glycans are prominently present in the profiles at low pH, but not at high pH (Supplemental Fig. 5).

Although endoS has been reported to be specific for IgG Fc, these experiments illustrate that this is only true at high pH. We speculated that the enzyme may recognize structurally similar protein substrates at suboptimal pH values. IgA and IgM are the two other immunoglobulins that are present in significant amounts in serum at about 2.5 µg/µl (IgA) and 1.5 µg/µl (IgM) (31). Moreover, the observed glycan types are typically abundantly present in IgA glycan profiles (32), but only at low levels in IgG profiles. Therefore, we assessed whether endoS can release IgA or IgM glycans at pH 5.0 and pH 8.5. We first confirmed the pH independence of IgG profiles and obtained identical profiles at pH 5.0 or 8.5 (Fig. 3B). EndoS digests at pH 5.0 of IgA purified from a fresh patient plasma sample (Fig. 3C, left) produced the same two sialylated non-IgG glycans as in IgG-depleted serum at pH 5.0. These glycans were absent in the profile of purified IgA digested at pH 8.5 (Fig. 3C, right). EndoS digests of commercial IgA gave the same result (data not shown). Similarly, endoS also produced minute amounts of the monosialylated N-glycan from IgM at pH 5.0 but not at pH 8.5 (data not shown).

To further investigate the contribution of IgA glycans, we compared the glycan profiles of fresh whole and IgA-depleted patient samples (Fig. 3D and 3E) at pH 5.0 and pH 8.5. Although completely removing IgA from the samples proved to be impossible even after several rounds of peptide M chromatography, a significant reduction in the peak height of the contaminating peaks in the pH 5.0 profile can be observed after IgA depletion. This indicates that these peaks are at least partially due to the presence of IgA. Taken together, these experiments indicate that the presence of other immunoglobulins does not interfere with the assay when working at a pH of 8.5 and that endoS is specific for IgG Fc glycans under these conditions.

As a final point, we also confirmed that, under the optimized assay conditions (pH 8.5), the presence or absence of serum components does not influence the undergalactosylation score in a cohort of patients and healthy controls with a varying degree of IgG undergalactosylation. To do this, we compared endoS derived profiles from serum versus purified IgG for the total patient cohort. An almost perfect correlation (Fig. 3F; Pearson’s r = 0.986, p < 0.001) and a regression line with slope 0.982 and intercept at 0.019 demonstrate that the presence of serum glycoproteins does not influence the outcome, underscoring the reliability of the assay in the selected conditions.

### EndoS enables direct serum IgG galactosylation assessment

Classically, IgG Fc galactosylation levels have been determined through PNGaseF mediated deglycosylation of purified whole IgG. There seems to be no general consensus on the choice of glycans for calculating the level of galactosylation. Often, the peak height or area of NGA2F (also known as G0F, sometimes just G0) is used, most often normalized to the total signal or to the monogalactosylated glycan peak NG1A2F (G1F, sometimes G1) (33,34). Another approach is to take the contribution of the galactosylation status of individual antennae on all glycans in the profile into account (35). In most cases, Fab fragment glycans also contribute to the calculation to some extent as PNGase is not specific for the Fc-linked N-glycan. On the other hand, under the conditions of our assay, endoS specifically cleaves N-glycans from IgG Fc fragments, while leaving Fab glycans intact. As a result, the measured undergalactosylation score (UGS) may not be exactly equal to that obtained from a PNGaseF digest. However, given that only 15-25% of Fab fragments are glycosylated, they contribute much less to the total IgG glycan content than do Fc fragments (100% glycosylated). In other words, Fab glycosylation is not expected to be a major confounder and the UGS calculated from endoS profiles should correlate well with scores calculated from PNGaseF-derived profiles. For the comparisons presented here, we decided to use the G0F level normalized to the total glycan signal for PNGaseF derived profiles, as this appears to be the most commonly used way of calculating galactosylation levels (Supplemental Fig. 2).

To assess the performance of our assay on patient serum samples, we analyzed IgG glycosylation in a total cohort of 188 patients and healthy controls with varying levels of galactosylation. We calculated the UGS from profiles obtained by either treating whole serum with endoS (and sialidase) or treating IgG purified from serum with PNGaseF and sialidase. We found a very good correlation between the two methods irrespective of the use of sialidase (Fig. 4A; no sialidase, Pearson’s r = 0.945, p < 0.001 and Fig. 4B; sialidase, Pearson’s r = 0.951, p < 0.001).

Data points that we identified as outliers in a residuals analysis (studentized residuals > 3) are indicated in red and represent the same samples in both experiments. Two outliers can be attributed to suboptimal IgG purification: the PNGaseF profiles from these samples contained relatively higher levels of triantennary glycan as compared to the rest of the cohort from which they were drawn. Triantennary glycans are not present on IgG and these glycans are thus considered to be an impurity from contaminating serum proteins. For these samples, it is likely that the UGS derived from PNGaseF profiles is incorrect. For the last outlier we found no straightforward explanation.

Taken together, these strong correlations show that the galactosylation state of IgG in serum can be reliably measured with endoS, without the need for IgG purification.

## Discussion

In current clinical practice, inflammation is predominantly determined by measuring CRP values in serum. However, the short half-life (18 hours) of CRP cause fluctuations on a daily basis. Therefore, CRP is only suitable to assess acute inflammation. For ESR, which is frequently used in routine clinical chemistry, the most restricting limitation is that it requires fresh blood. Moreover, it has several analytical interferences that may confound the results (14). Hence, there is an unmet need for a marker that measures cumulative exposure to inflammation.

Changes in IgG Fc glycosylation are uniquely suitable for monitoring chronic inflammation because IgG has a half-life of 24 days. The inflammatory processes that lead to IgG glycosylation changes need to persist for at least one to two t1/2 of the IgG pool to become noticeable. Since the first report in the context of RA in 1985 (1), many publications reported a decrease in IgG galactosylation in the vast majority of chronic necro-inflammatory diseases (2–9) and recently very large studies have been reported on this glycosylation change (36,37). So far, all assays require IgG purification followed by complex analytics, making the biomarker rather unsuitable for routine clinical chemistry.

Here we used endoglycosidase S from a pathogenic bacterium to cleave the N-glycans of IgG Fc. We show that, under optimized conditions, this enzyme is sufficiently specific that it can be used in a highly complex sample such as serum. We found that the enzyme’s specificity depends on the pH of the reaction buffer. In the pH range of 5.0-8.0, endoS is not entirely IgG specific, as it also releases some IgA N-glycans and minute amounts of IgM N-glycans. The specificity of endoS for IgG Fc drops with increasingly lower pH values of the buffer. However, under optimal conditions (at pH 8.5 or higher) we only detected glycans that were derived from IgG Fc fragments.

Whole IgG UGS measurements by PNGaseF strongly rely on the purity of the IgG, which inherently is a potential problem during sample preparation. Moreover, since Fab and Fc glycans are distinct but overlapping, contribution of Fab N-glycans can never be completely excluded as a confounding factor, even when using purified IgG. Because of its specificity for IgG Fc under the assay conditions, UGS assessed by endoS should be more reliable than UGS from PNGaseF profiles of purified IgG.

By comparing endoS profiles of serum and purified IgG in a cohort of patients and healthy controls, we confirmed that the outcome of the endoS-based assay is not dependent on the presence of other glycoproteins: we found an almost identical outcome for both methods.

When employing CE as an analysis method, glycans are often treated with sialidase to obtain a profile that is easier to interpret and quantify. However, the endoS-derived glycan profile is quite simple, rendering the use of sialidase for proper peak identification and quantification unnecessary. We also showed that sialidase treatment of endoS-released glycans does not change the outcome of our assay and therefore propose to omit it in the final protocol. This results in an even faster and cheaper workflow requiring fewer analyte-specific reagents.

Our new endoS-based assay thus represents an important breakthrough towards the clinical implementation of the well-studied IgG undergalactosylation as a correlate of chronic inflammation. It encompasses a fast and simple, Fc-glycan specific sample preparation and subsequent analysis on high-throughput CE analyzers, e.g. on capillary DNA sequencers. The analytical step can also be readily adapted to cheap CE-based microfluidics platforms as we have shown before (25) and indeed clinical CE analysers (38), paving the road for implementation in clinical chemistry routine. We envisage that this new assay will find use in diagnosis of the presence of chronic inflammation (e.g. as it accompanies the senescence processes associated with old age) and in monitoring of responses to anti-inflammatory drug treatments.

## Abbreviations

APTS: 8-Aminopyrene-1,3,6-trisulfonic acid
CE: capillary electrophoresis
CE-LIF: Capillary electrophoresis – laser-induced fluorescence
CRP: C-reactive protein
CV: column volume
EndoS: endo-β-N-acetyl-glucosaminidase S from *Streptococcus pyogenes* = Endoglycosidase S
ESR: erythrocyte sedimentation rate
GlcNAc: N-acetylglucosamine
gp120: HIV envelope glycoprotein gp120
HBV: Hepatitis B virus
MS: mass spectrometry
NAFLD: non-alcoholic fatty liver disease
NASH: non-alcoholic steatohepatitis
PNGaseF: Peptide N-glycosidase from *Elizabethkingia meningoseptica* (previously *Flavobacterium meningosepticum)*
RA: rheumatoid arthritis
TNF: tumor necrosis factor
UGS: UnderGalactosylation Score
UPLC: ultra-performance liquid chromatography

## Acknowledgements

The research in this manuscript was supported by an ERC grant (ERC-2013-CoG-616966).

