## Supplementary Materials for "Endoglycosidase S enables a highly simplified clinical chemistry assay procedure for direct assessment of serum IgG undergalactosylation in chronic inflammatory disease"

### Supplemental data

#### Contents

Supplemental data ... 1
- Supp. Fig. 1: symbolic glycan representation and nomenclature ... 2
- Supp. Fig. 2: Formulae ... 3
- Supp. Fig. 3: exoglycosidase digests of endoS-derived IgG glycans ... 4
- Supp. Fig. 4: endoS sensitive and resistant N-glycans on IgG Fc and Fab ... 5
- Supp. Fig. 5: pH dependence of IgG-depleted serum and repleted serum ... 7
- References ... 7

#### Supp. Fig. 1: symbolic glycan representation and nomenclature

**PNGase F-released glycans, symbols, abbreviations and condensed nomenclature**

| | | |
|---|---|---|
| 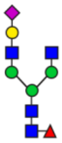 | N1G1A2F | Neu(α-2,6)Gal(β-1,4)GlcNAc(β-1,2)Man(α-1,3)[GlcNAc(β-1,2)Man(α-1,6)] Man(β-1,4)GlcNAc(β-1,4)[Fuc(α-1,6)]GlcNAc |
| 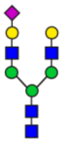 | N1A2 | Neu(α-2,6)Gal(β-1,4)GlcNAc(β-1,2)Man(α-1,3)[Gal(β-1,4)GlcNAc(β-1,2) Man(α-1,6)]Man(β-1,4)GlcNAc(β-1,4)GlcNAc |
| 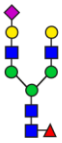 | N1A2F | Neu(α-2,6)Gal(β-1,4)GlcNAc(β-1,2)Man(α-1,3)[Gal(β-1,4)GlcNAc(β-1,2) Man(α-1,6)]Man(β-1,4)GlcNAc(β-1,4)[Fuc(α-1,6)]GlcNAc |
| 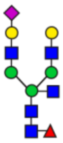 | N1A2FB | Neu(α-2,6)Gal(β-1,4)GlcNAc(β-1,2)Man(α-1,3)[GlcNAc(β-1,4)][Gal(β-1,4) GlcNAc(β-1,2)Man(α-1,6)]Man(β-1,4)GlcNAc(β-1,4)[Fuc(α-1,6)]GlcNAc |
| 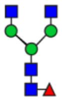 | NGA2F | GlcNAc(β-1,2)Man(α-1,3)[GlcNAc(β-1,2)Man(α-1,6)] Man(β-1,4)GlcNAc(β-1,4)[Fuc(α-1,6)]GlcNAc |
| 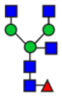 | NGA2FB | GlcNAc(β-1,2)Man(α-1,3)[GlcNAc(β-1,4)][GlcNAc(β-1,2)Man(α-1,6)] Man(β-1,4)GlcNAc(β-1,4)[Fuc(α-1,6)]GlcNAc |
| 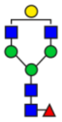 | NG1A2F | $NG1A2F_1$ = Gal(β-1,4)GlcNAc(β-1,2)Man(α-1,3)[GlcNAc(β-1,2)Man(α-1,6)] Man(β-1,4)GlcNAc(β-1,4)[Fuc(α-1,6)]GlcNAc<br>Or<br>$NG1A2F_2$ = GlcNAc(β-1,2)Man(α-1,3)[Gal(β-1,4)GlcNAc(β-1,2)Man(α-1,6)] Man(β-1,4)GlcNAc(β-1,4)[Fuc(α-1,6)]GlcNAc |
| 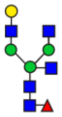 | NG1A2FB | Gal(β-1,4)GlcNAc(β-1,2)Man(α-1,3)[GlcNAc(β-1,4)][Gal(β-1,4) GlcNAc(β-1,2)Man(α-1,6)]Man(β-1,4)GlcNAc(β-1,4)[Fuc(α-1,6)]GlcNAc |
| 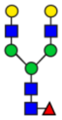 | NA2F | Gal(β-1,4)GlcNAc(β-1,2)Man(α-1,3)[Gal(β-1,4)GlcNAc(β-1,2)Man(α-1,6)] Man(β-1,4)GlcNAc(β-1,4)[Fuc(α-1,6)]GlcNAc |
| 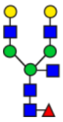 | NA2FB | Gal(β-1,4)GlcNAc(β-1,2)Man(α-1,3)[GlcNAc(β-1,4)][Gal(β-1,4) GlcNAc(β-1,2)Man(α-1,6)]Man(β-1,4)GlcNAc(β-1,4)[Fuc(α-1,6)]GlcNAc |

**EndoS-released glycans, symbols, abbreviations and condensed nomenclature:**

| | | |
|---|---|---|
| 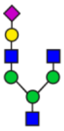 | N1G1A2F* | Neu(α-2,6)Gal(β-1,4)GlcNAc(β-1,2)Man(α-1,3)[GlcNAc(β-1,2)Man(α-1,6)] Man(β-1,4)GlcNAc |
| 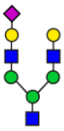 | N1A2F* | Neu(α-2,6)Gal(β-1,4)GlcNAc(β-1,2)Man(α-1,3)[Gal(β-1,4)GlcNAc(β-1,2) Man(α-1,6)]Man(β-1,4)GlcNAc |
| 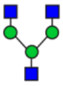 | NGA2F* | GlcNAc(β-1,2)Man(α-1,3)[GlcNAc(β-1,2)Man(α-1,6)] Man(β-1,4)GlcNAc |
| 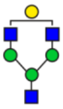 | NG1A2F* | $NG1A2F_1^*$ = Gal(β-1,4)GlcNAc(β-1,2)Man(α-1,3)[GlcNAc(β-1,2)Man(α-1,6)] Man(β-1,4)GlcNAc<br>Or<br>$NG1A2F_2^*$ = GlcNAc(β-1,2)Man(α-1,3)[Gal(β-1,4)GlcNAc(β-1,2)Man(α-1,6)] Man(β-1,4)GlcNAc |
| 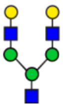 | NA2F* | Gal(β-1,4)GlcNAc(β-1,2)Man(α-1,3)[Gal(β-1,4)GlcNAc(β-1,2)Man(α-1,6)] Man(β-1,4)GlcNAc |

* The first GlcNAc and core fucose are not present on N-glycan structures that are cleaved by endoS. Symbolic representation and color coding for the monosaccharides follows the guidelines of the consortium for functional glycomics (1).

#### Supp. Fig. 2: Formulae

$$\boldsymbol{PNGase\ UGS} = \frac{NGA2F}{NGA2 + NGA2F + NGA2FB + NG1A2F_1 + NG1A2F_2 + NA2 + NA2F + NA2FB}$$

$$\boldsymbol{endoS\ UGS} = \frac{(2 * (NGA2F^*) + (NG1A2F_1^* + NG1A2F_2^*))}{\left(2 * (NGA2F^* + NG1A2F_1^* + NG1A2F_2^* + NA2F_1^*)\right)}$$

**Supplemental Figure 2:** Formulae used to calculate UGS from endoS and PNGaseF profiles. All formulae use the shorthand notation for each glycan as found in Supplemental figure 1.

Supp. Fig. 3: exoglycosidase digests of endoS-derived IgG glycans

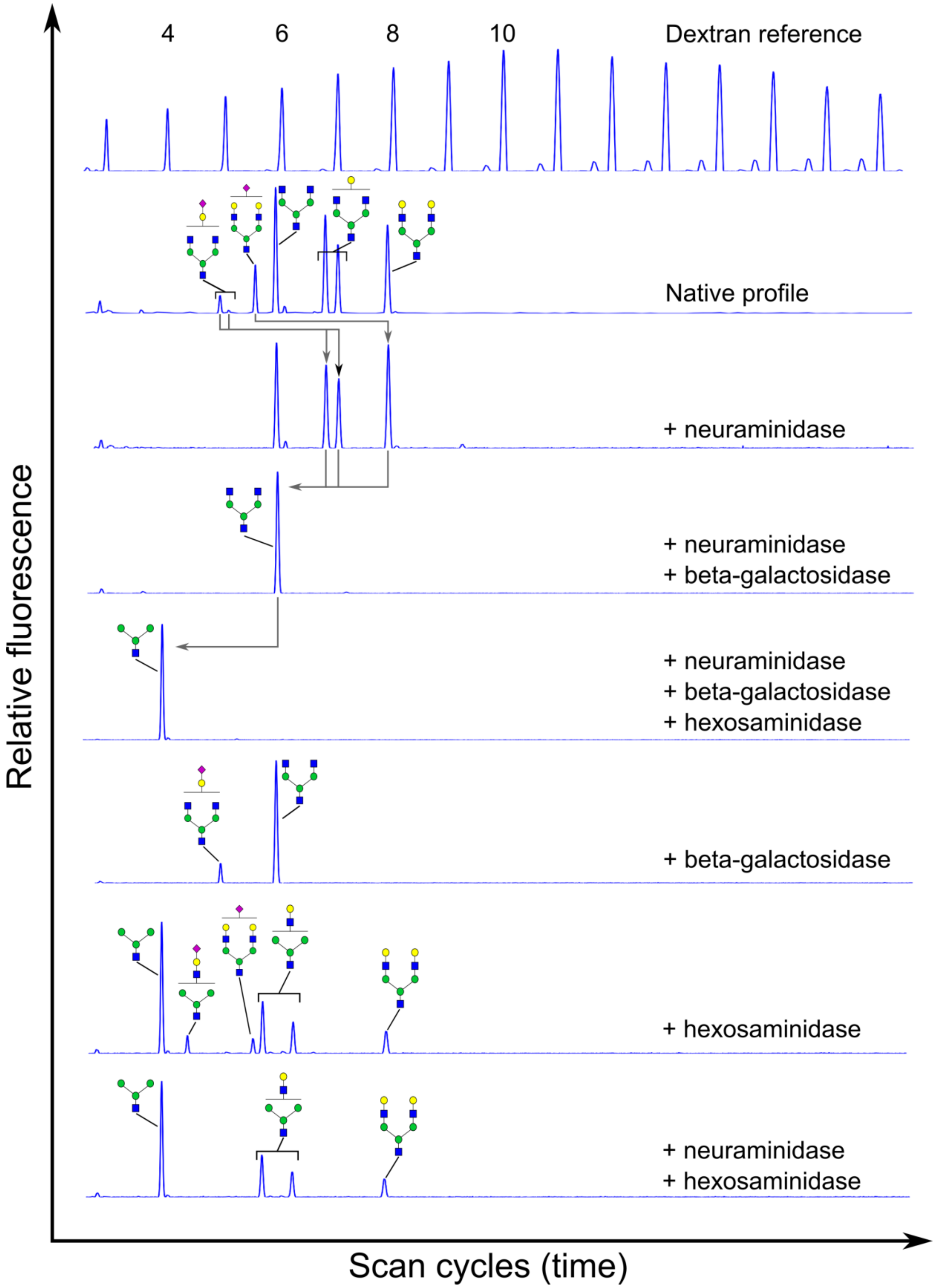

**Supplemental Figure 3**: Serum-derived N-glycans after endoS treatment (at pH 8.5) were labelled and analysed by CE-LIF. Various exoglycosidase digests were performed: *A. ureafaciens* neuraminidase (sialidase, in house production), *S. pneumonia* β-1,4-galactosidase (Prozyme), Jack bean hexosaminidase (Prozyme) or combinations thereof, revealing the structure of the endoS-released N-glycans, as indicated in the panels. In the top panel, a dextran reference ladder (for neutral N-glycans) is shown, and the number of monosaccharides is indicated. Note that sialylated N-glycans run below their expected size, since they carry an extra negative charge and thus migrate faster through the capillary than do neutral N-glycans.

**Supplemental Figure 3 Methods:** Methods are the same as in the main article for endoS and as previously described (2) for exoglycosidase digests. Briefly, labelled glycans were dried, reconstituted in an appropriate buffer and treated overnight at 37°C with exoglycosidases.

#### Supp. Fig. 4: endoS sensitive and resistant N-glycans on IgG Fc and Fab

To assess which IgG Fc N-glycans are hydrolysed by endoS and to verify Fab N-glycans are insensitive to endoS, we compared the glycans on Fab and Fc fragments with or without endoS pre-treatment. To do this, whole IgG was either treated with endoS or not, then bound to protein A magnetic beads and extensively washed. After the washing steps, IgG was digested with IdeS (FabRICATOR, Genovis) on the beads to release $F(ab)_2$ fragments. These were collected and the beads were again extensively washed, after which the Fc fragments were eluted and collected.

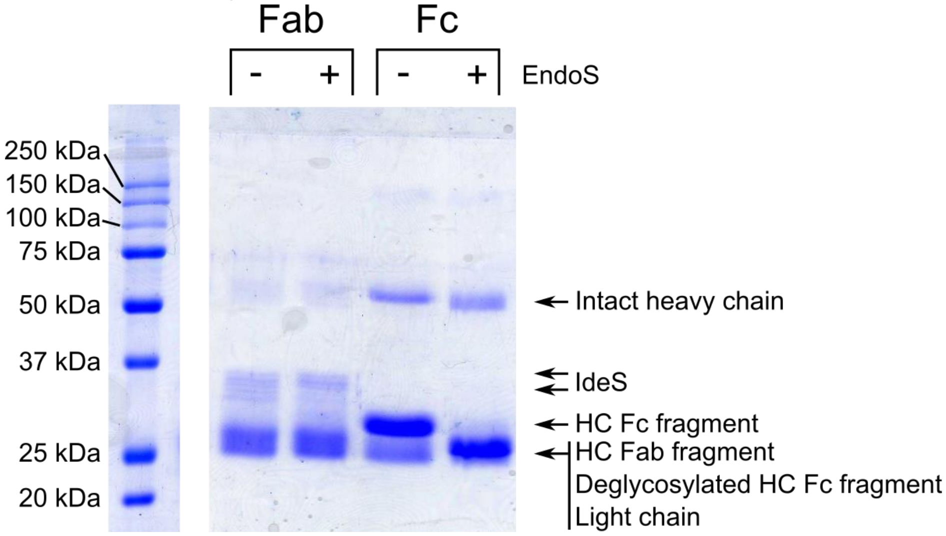


**Supplemental Figure 4a**: A Coomassie-stained reducing SDS-PAGE gel shows each fraction, allowing to assess IdeS digest completeness and fragment purity. A significant enrichment of Fc and Fab fragments in each fraction is obtained; the heavy chain Fc band cannot be observed in the Fab fractions (lane 1 and 2), while a significant excess of heavy chain Fc fragment over Fab fragment constituents is observed in lanes 3 and 4. However, even after extensive IdeS digestion (several aliquots of excess IdeS, combined with overnight digestion), some intact IgG heavy chain remains in the Fc fractions as can be seen in the Fc lanes (lanes 3 and 4).

A shift in MW of several kDA is observed for the HC Fc fragment after endoS hydrolysis (lane 3 and 4), while no difference is observed in the light chain and heavy chain Fab fragments (lane 1 and 2), already hinting to the Fc-specificity of endoS.

**Supplemental Figure 4b**: Glycans of the Fab and Fc fractions were released with PNGaseF from both fraction of both the endoS- and buffer-treated samples. The first trace shows a whole IgG PNGaseF digest for comparison, with peaks annotated. The second and fourth traces show Fc and Fab glycans without endoS pre-treatment, revealing mainly bisialylated and some monosialylated biantennary N-glycans as the most Fab N-glycans. All other glycans are almost exclusively found on Fc. Minor amounts of the bisialylated N-glycans are observed in the Fc trace. These are most likely Fab glycans in that fraction, due to an incomplete IdeS digest.
Comparing both Fc traces, it becomes clear that the glycans carrying a bisecting N-Acetylglucosamine are resistant to endoS, which is in agreement with previously published results (3). All other Fc glycans are fully hydrolysed by endoS.

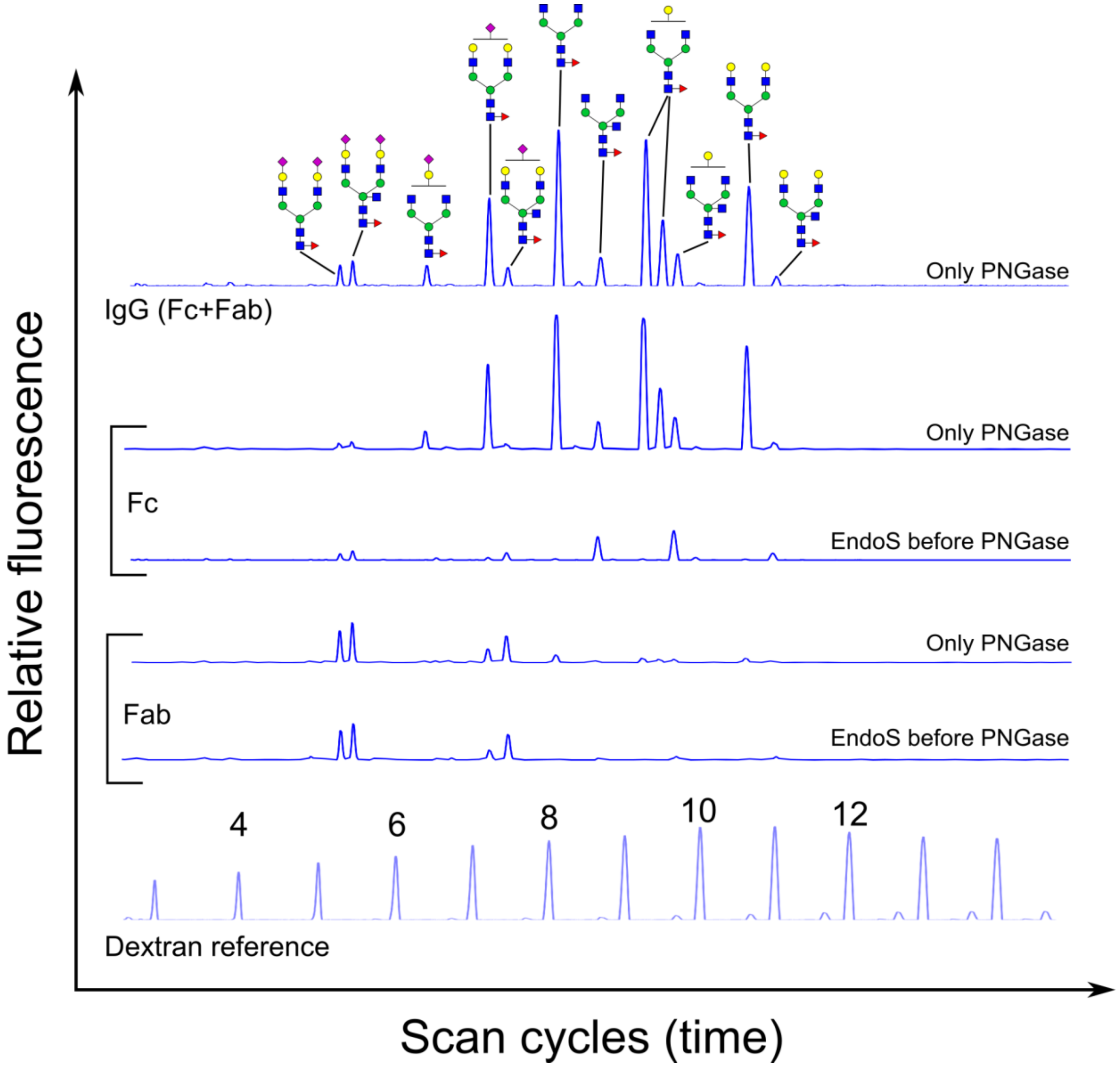


**Supplemental Figure 4 methods:** 100 µg of IgG (IgG purified from human serum, purity > 95%, Sigma-Aldrich) was dissolved at 10 µg/µl in PBS and was treated for 3 hours at 37°C with 100 units of endoS (Genovis) in 50 mM Tris-Cl + 150 mM NaCl, pH 8.5. A second identical sample was incubated in buffer alone. Both samples were then loaded on protein A magnetic beads (New England Biolabs), as per the manufacurer's instructions. After binding, the beads were extensively washed with binding buffer. Then, 150 units of IdeS (FabRICATOR, Genovis) were added to the beads in 10 mM sodium phosphate buffer pH 8.0, the samples were incubated for 3 hours at 37°C, and then a second aliquot of 150 units of IdeS was added and the samples were incubated at 37°C overnight. The following morning, the supernatants containing the released $F(ab)_2$ fragments were collected. The beads were again extensively washed

with binding buffer and finally the Fc fragments were eluted with 200 mM glycine buffer pH 2.5. The Fc fragments were then neutralized with an equal volume of 1 M Tric-Cl, pH 9.0.
The Fc and F(ab)$_2$ fragments from endoS-treated and control IgG were then used as the starting material for on membrane N-glycan preparation as described previously (2).

#### Supp. Fig. 5: pH dependence of IgG-depleted serum and repleted serum

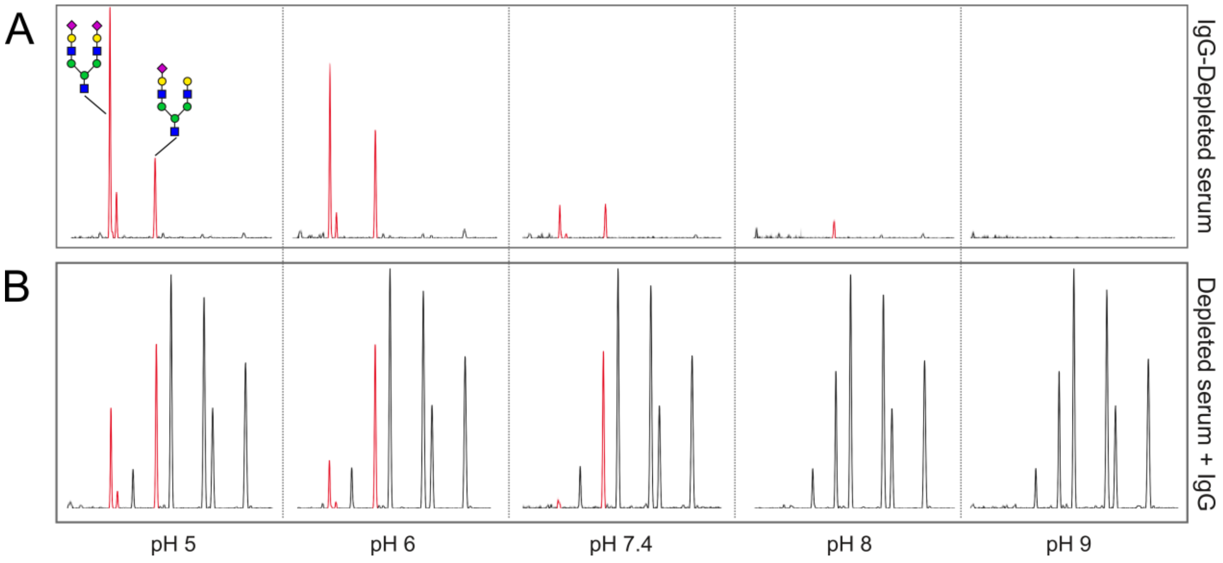


**Supplemental Figure 5:** Panel A is the same as Figure 3A in the main article. It shows that treating IgG-depleted fresh serum with endoS results in the release of non-IgG N-glycans at low pH. Adding a physiological amount (10 µg/µl) of commercial IgG back to the depleted serum shows that non-IgG N-glycans are prominent in the profile at low pH, resulting in a profile that is different from the profile obtained from purified IgG (see main article). These contaminating peaks are absent at pH 8.0 or higher, resulting in a profile that is indistinguishable from that of pure IgG.

Methods for this figure are as described in the main article.
